# Nutritional characterisation and seasonal variation of goat forages in Central Malawi

**DOI:** 10.1101/2023.12.22.573017

**Authors:** A. S. Cooke, W. Mvula, P. Nalivata, J. Ventura-Cordero, L.C. Gwiriri, T. Takahashi, E. R. Morgan, M. R. F. Lee, A. Safalaoh

## Abstract

Goat ownership is prevalent across rural Malawi and provides a vital source of income, nutrition, and food security. However, goat performance is poor, and this presents a risk to individuals and communities who depend on them. Whilst mitigating this through supplementation of persevered forages may be possible, this is a challenge due to limitations of resources and the fact that goats typically free-roam during the dry season. Nutrition is fundamental to health and productivity of any livestock enterprise, it is required to be able to deal with stresses, such as disease, and to enable growth and production. The aim of this study was to characterise the nutritional profile of naturally available forages in Malawi, including the seasonal variation in nutrition. Samples of herbaceous forages and browse were collected over a 17-month period, across four villages (30 farms/smallholders) in Central Malawi. Forages underwent nutritional analysis for crude protein, fibre fractions, and ash/organic matter and NDVI obtained from satellite imagery was used as a measure of forage availability. Forage nutrition and availability were most adequate in the wet season, with higher concentrations of crude protein and a greater availability of herbaceous plants. There were significant differences in low-digestibility fibre fractions between locations, likely due to local factors such as soil and hydrology. The fall in crude protein concentrations from the wet season to the dry season represent a seasonal nutrition-gap which may result in risks to goat health and productivity.

## 1 Introduction

Goat ownership is common across Sub-Saharan Africa with household ownership levels especially high in rural communities (Gwiriri et al., 2023; Kaumbata et al., 2020; Taruvinga et al., 2022). The national goat herd of Malawi is estimated to be in excess of 10 million head, increasing rapidly, from 3.9 million in 2010 and 1.7 million in 2000 (FAO, 2022). Individual herd sizes are typically small with a mean of 8 head (Kaumbata et al., 2020). Their meat and milk provide high density nutrition to the human population, which is pertinent given the high incidence of malnutrition and food insecurity in the country (NSO, 2017). Goats can also act as a form income, through the direct sale of meat, milk, and live animals, and as a form of insurance, a store of economic and nutritional resources that can be utilised in times of difficulty (Malata and Banda, 2009). Kaumbata et al. (2020) calculated that goat ownership yielded a 25% net return, rising to 39% when considering intangible benefits. The study found that live sales comprised the majority (79%) of off-take, suggesting that goats are predominantly reared for their cash value. Goats typically sold for MK15,000 to 20,000 (Malawian Kwacha, approx. USD$18.50 to $25.00). Malata and Banda (2009) reported that females were most sold (75% of sales), especially breeding females (49%). Goat sales in Malawi typically function on a per head basis (as opposed to per kg liveweight as common elsewhere). The combination of this sales model and the relatively high turnover of animals means that goat mortality is a significant risk to income and food security in both the short and long term. Despite this, mortality is high with rates of 3-10% for adult goats and 8-26% for kids, though anecdotally rates can be far higher for individual smallholders. Whilst predation is the largest cause of mortality (25%), diarrhoea (19%), disease (11%), and starvation (8%) are also major causes (Kaumbata et al., 2020).

Mortality and productivity issues, as outlined by Kaumbata et al. (2020), can partially be mitigated through nutritional interventions. Nutrition is fundamental to goat health and productivity. Improving nutrition can reduce susceptibility to disease, increase reproductive success, and provide greater yields of meat and milk, all of which yield socio-economic benefits to owners (Kaumbata et al., 2020; NRC, 2006). For example, Ajayi et al, (2008) found feed interventions to significantly increase the performance of West African Dwarf goats, though there has been little conducted exploring this in Malawi. Whilst enhancing nutrition provides benefit, inadequate nutrition can be deleterious and was identified by Bath et al. (2005) as one of the four major health/disease categories for goats in southern Africa. Whilst supplementation and other interventions can be effective in improving nutrition, they are often unavailable or too costly for smallholders in the region. Therefore, optimising the use of local and naturally available forages should be the first step in any nutritional intervention strategy.

Malawi has distinct wet and dry seasons (Figure 1). During wet seasons temperatures are high, typically reaching a monthly average of 25.7°C in November with rainfall peaking at 243 mm in January. The dry season is cooler with a monthly mean temperature as low as 18.7°C in July and rainfall of < 3 mm in September (The World Bank, 2021). This seasonality inevitably impacts forage availability and nutrition, which is a significant issue across the region (Cooke et al., 2023b). Becker and Lohrmann (1992) studied goat forage nutrition and selection in Malawi and found significant seasonal differences in forage nutrition. From the end of the wet season (March to June) to the end of the dry season (September to October), crude protein and hemicellulose concentrations typically decreased whilst cellulose concentrations increased. Similar seasonal variation in forage has been observed elsewhere across Sub-Saharan Africa (Naumann et al., 2017; Omphile et al., 2005; Setshogo et al., 2011).

**Figure 1.**
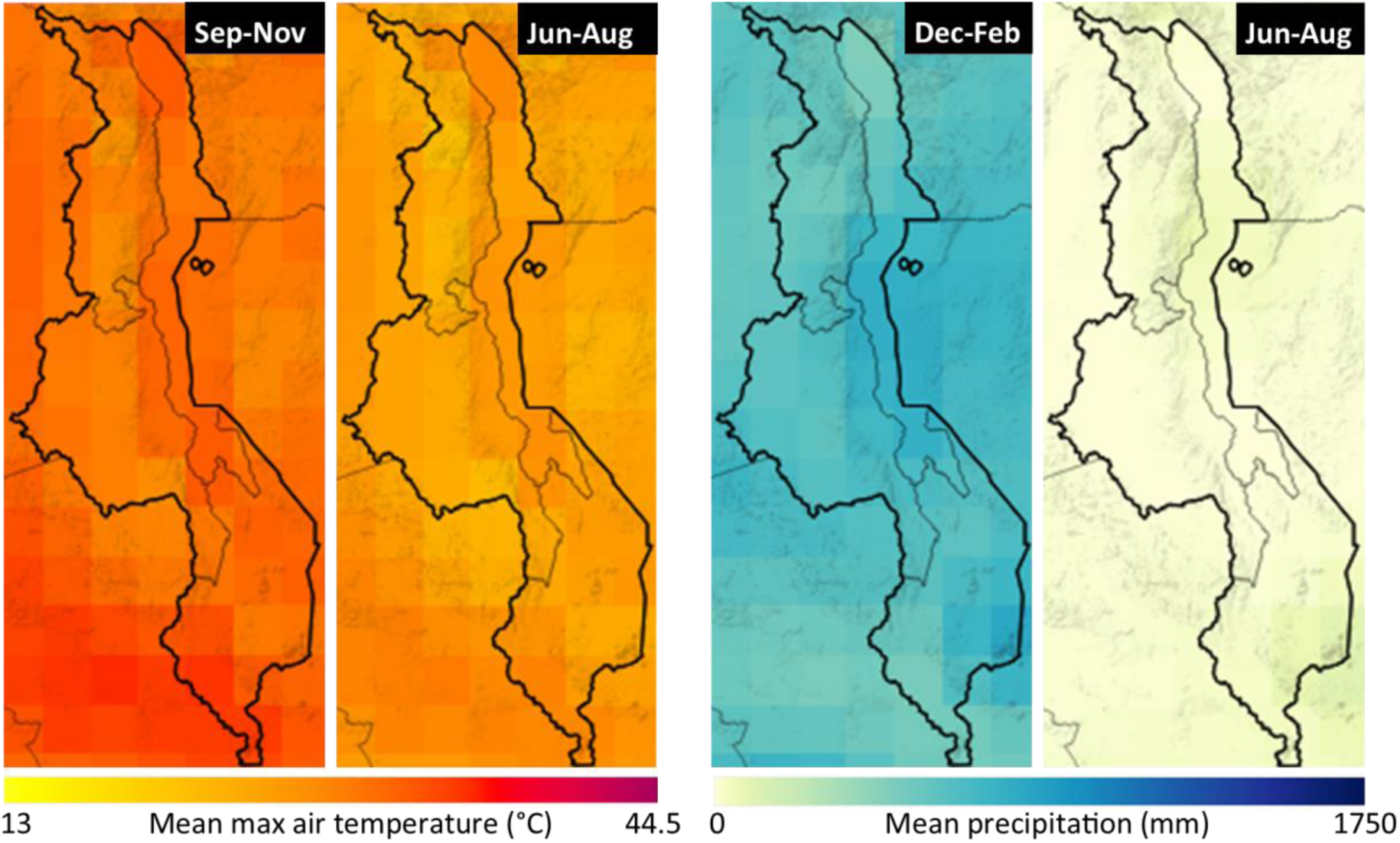
Mean maximum air temperature (°C) and mean total precipitation (mm) for the most extreme months of the wet season (left of pair) and dry season (right of pair). Data is a mean from 1991-2020. Data and images taken from The World Bank (2021) and altered under license CC BY 4.0.

Goat husbandry is an essential factor to account for when considering this issue. Typically, during the dry season, once crops have been harvested, goats graze freely. During the wet season, to protect crops, goats are tethered or kept in communal areas (Banda et al., 1993; Kaumbata et al., 2020). This provides both opportunity and challenge. Opportunity in the wet season to utilise available nutrition through husbandry/housing practices, and challenges in the dry season to provide nutritional intervention when goats are free grazing. Finally, as the pressures of climate change increase, disproportionately affecting the global south, we can expect to see the extremes of seasonality and increase along with the issues it can cause (Dunning et al., 2018; Lawal et al., 2019).

The objective of this study was to characterise the nutritional profile of naturally available forages in Malawi. Seasonal variation in forage nutrition was assessed and combined with satellite data on forage abundance to provide a holistic perspective seasonal nutritional quality and quantity. Finally, results were considered with respect to each of the study sites.

## 2 Methods

The study was conducted in the Central Region of Malawi, across four villages approximately 20-25 km south of the capital city, Lilongwe. Annual rainfall is typically in the region of 800-1000 mm and the average annual temperature in the region of 23-25°C (NASA, 2022a). However, there are strong seasonal variations in weather with little to no rainfall and temperatures in the low teens in the dry season. The study area was on lixisol, which is a heavily weathered clay soil with low nutrient levels (ISRIC, 2006).

Forage samples (136 samples, 26 species) were collected between 02/2020 and 07/2021 in the Central Region of Malawi across four villages, covering 30 smallholders: Chinkhowe (*n* = 50, 7 farms), Kamchedzera (*n* =23, 12 farms), Mazinga (*n* = 32, 6 farms), and Mkwinda (*n* = 31, 5 farms) (Figure 2). Plants were chosen for sampling based on informal observation of goat foraging and browsing behaviour and on farmer discussions. Broadly, plant material came from within pens, near tethering points, or from grazing land. Samples from pens included the mixed grasses and forages provided by farmers to the goats. Samples near tethering points were taken within a 4 m radius and based on a 30 min observation of what goats were grazing/browsing. Samples from grazing areas were also chosen based on 30 min observations. To the greatest extent practicable, sampling excluded woody stems, clearly diseased or dead plant material, and seeds. Roots or any foreign material attached to the sample was removed and discarded. For grasses, leaf blades were clipped by hand to avoid contamination with soil or other material. Due to the seasonality of different plants and temporal timescale of sampling, plants were at different growth stages at collection, though most leaves samples were fully developed. Each sample was identified to genus or species level and categorised as either ‘herbaceous’ (soft stemmed plants such as grasses and forbs, *n* = 55) or ‘browse’ (hard stemmed plants such as trees, bushes, and shrubs, *n* = 81). Collection location and date were recorded, and samples classified as being collected in either the wet season (November to April, *n* = 76) or dry season (May to October, *n* = 60).

**Figure 2.**
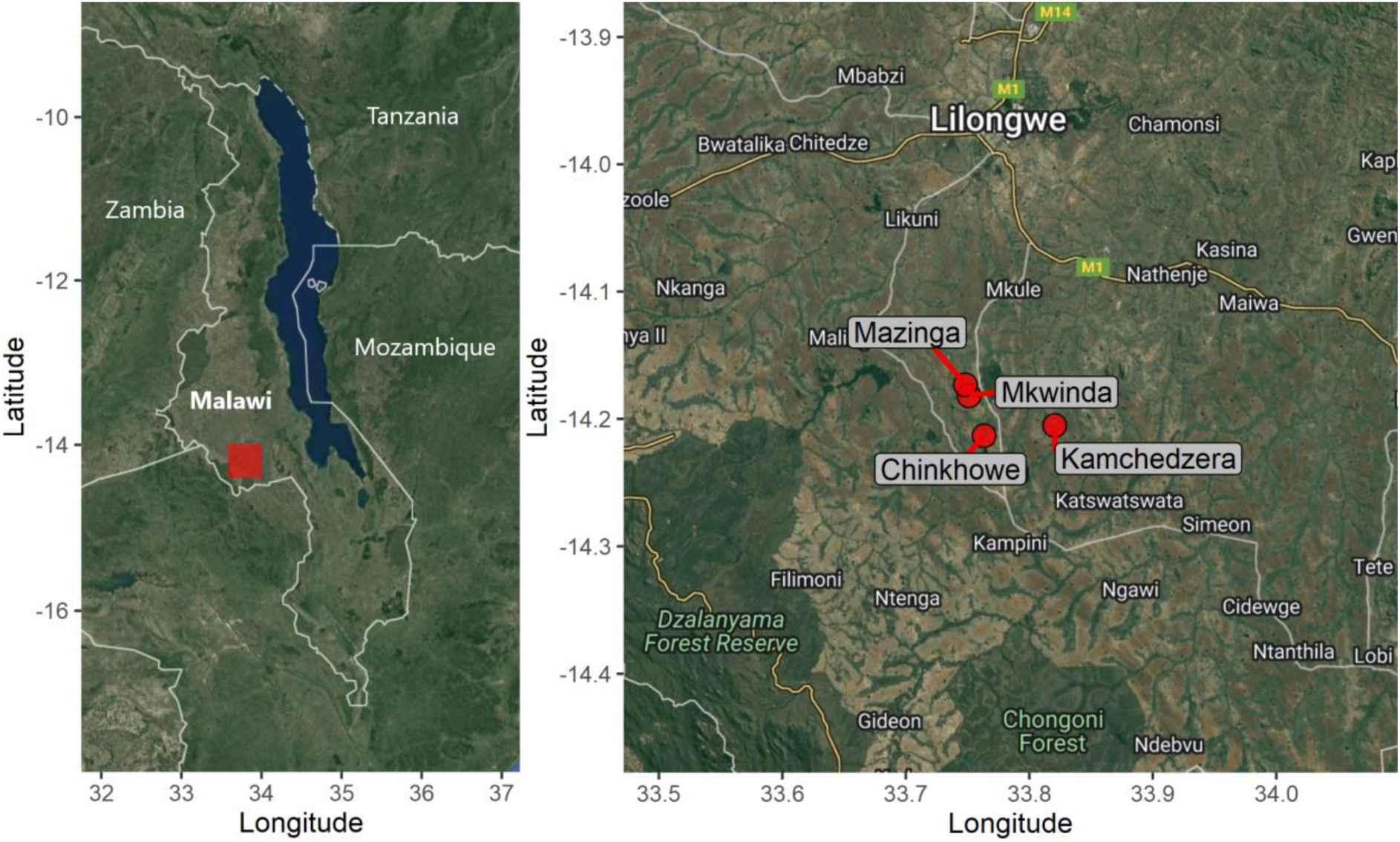
Map of study areas. Left: Map of Malawi with approximate study area in red. Right: Closeup of study area with site locations identified. Base map taken from Google maps (Google, 2021).

Samples were oven dried to a constant weight in Malawi before being vacuum packed and shipped to the UK where they were freeze dried and ground to < 2 mm. Loss on ignition was conducted (0.5 g, 540°C, 6 h) to determine organic matter (OM) and subsequently ash content by mass difference. Crude protein (CP) was determined as 6.25x nitrogen content, as determined by the Dumas technique. Four fibre fractions were determined using an ANKOM 200 Fiber Analyzer based on methods by Goering and Soest (1970): neutral detergent fibre (NDF), acid detergent fibre (ADF), and acid detergent lignin (ADL). Hemicellulose was determined as NDF less ADF and cellulose as ADF less ADL.

### 2.1 Data analysis

An ANOVA was conducted to assess the relationships of forage species, forage type (herbaceous or browse), location (village), and season. Principal component analysis (PCA) was used to present the variation in forage nutrition relative to a hypothetical ‘ideal’, realistic, composition of ash (9%), protein (15%), lignin (8%), hemicellulose (18%), and cellulose (18%). These parameters were set based off of the literature (Lu et al., 2005; NRC, 2006; Pugh, 2014; Salah et al., 2014).

Additional figures were also created to represent the seasonal fluctuation in forage availability. ). NDVI data was used as a proxy for the overall availability of healthy vegetation available at each site. NDVI has been found to correlated reasonably well with biomass (Butterfield and Malmström, 2009; Calvão and Palmeirim, 2004) and notably, Wessels et al. (2006) found a correlation of R^2^ = 0.76 across different landscapes in Kruger National Park, South Africa where a landscape is comparable to the study area.Firstly, 16-day NDVI data was taken at a 250m^2^ resolution from the Moderate Resolution Imaging Spectroradiometer (MODIS) aboard the Terra satellite (NASA, 2022b). This was plotted around the study areas for both the heights of the dry season (specifically 12/08/2020 to 27/08/2020) and wet season (02/02/2021 to 17/02/2021). Secondly, mean CP values were calculated for all browse and all herbaceous species on a monthly basis. Monthly NDVI data was taken from MODIS/Terra on a 1km^2^ and the mean NDVI calculated across the four sites for each month. This was plotted against the monthly CP data. Statistical analysis and graphing was performed using R (version: 4.22) and R Studio (version: 2022.07.02) (R Core Team, 2021) (R Studio Team, 2020) and mapping in QGIS (version 3.28.1) (QGIS, 2022).

## 3 Results

Forage nutrition varied significantly between forage type, location, and season (Table 1, Table 2). Both OM/ash and NDF varied significantly with forage type, with marginally higher levels of OM in browse species compared to herbaceous (88.9% vs. 87.5%) but far lower concentrations of NDF (39.8% vs 54.3%) (Table 1). There was a significant difference in ADF, ADL, and between locations with plants sampled from Kamchedzera having the highest concentrations of ADF and ADL and Chinkhowe the lowest. There was a significant effect of season on CP and ADF, with higher CP concentrations observed in the wet season (14.4% vs 17.1%) and also higher ADF (28.1% vs 28.7%).

**Table 1.**
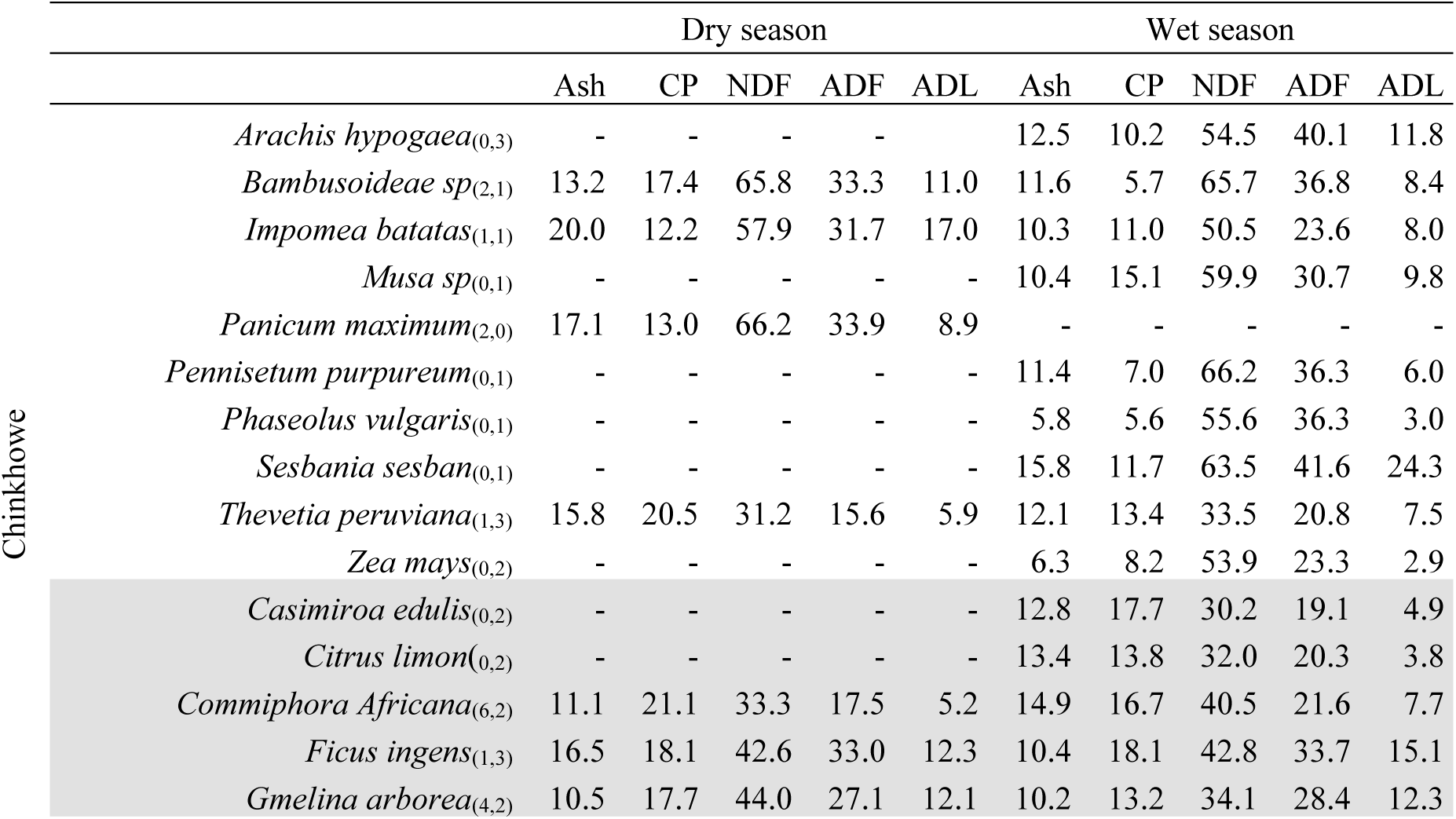
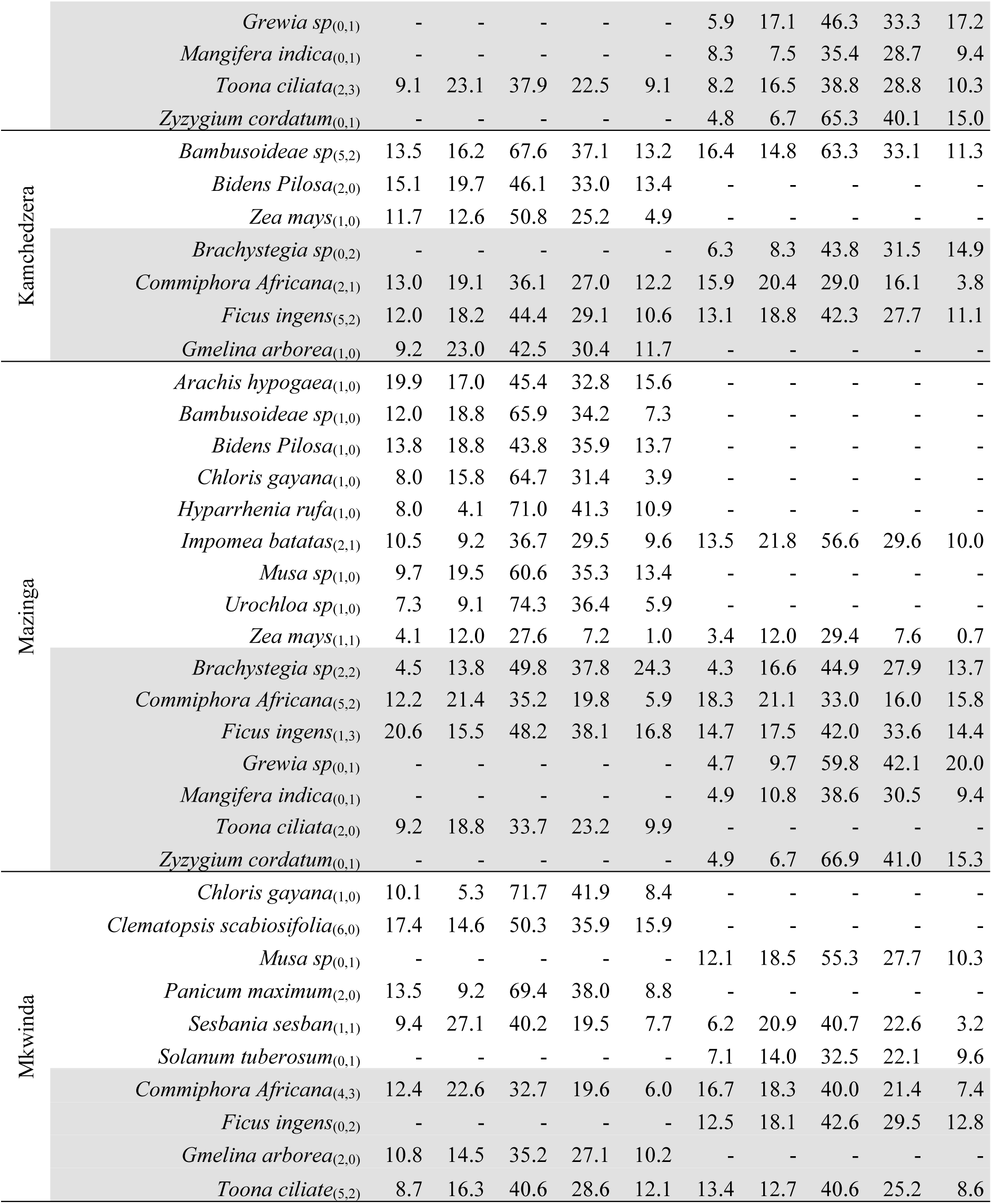
Nutritional values of all plant species split by location and season. Plant species not shaded in grey are ‘herbaceous’, plant species shaded in grey are ‘browse’. Values are means. - signifies that no sample was available/collected. Subscript after species names represents sample number (*n*) for the wet and dry season respectively.

**Table 2.**
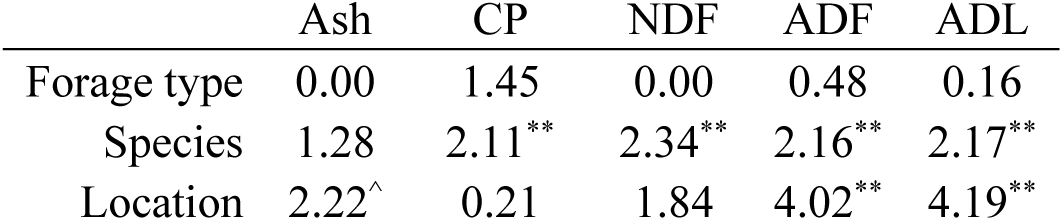
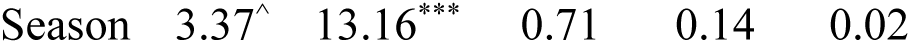
ANOVA results (*F-*values) comparing the concentrations of Ash, CP, NDF, ADF and ADL based on explanatory variables of forage type (herbaceous or browse), species, location, and season. Superscript symbols represent p-values: *** 0.001 ** 0.01 * 0.05 ^ 0.10.

|  | Ash | CP | NDF | ADF | ADL |
| --- | --- | --- | --- | --- | --- |
| Forage type | 0.00 | 1.45 | 0.00 | 0.48 | 0.16 |
| Species | 1.28 | 2.11** | 2.34** | 2.16** | 2.17** |
| Location | 2.22^ | 0.21 | 1.84 | 4.02** | 4.19** |

The first principal component (PC1) accounted for 40.0% of variation whilst the second (PC2) accounted for 22.3% (Figure 3). The PCA biplot shows how samples from Mazinga had a high degree of variation, especially compared to those from Kamchedzera and Mkwinda, which were more centrally concentrated on both PC1 (40.0% of variance) and PC2 (22.3% of variance). Seasonal variation is also apparent, with plants obtained during the wet season being more towards the negative end of PC1, which most strongly correlates with increased CP.

**Figure 3.**
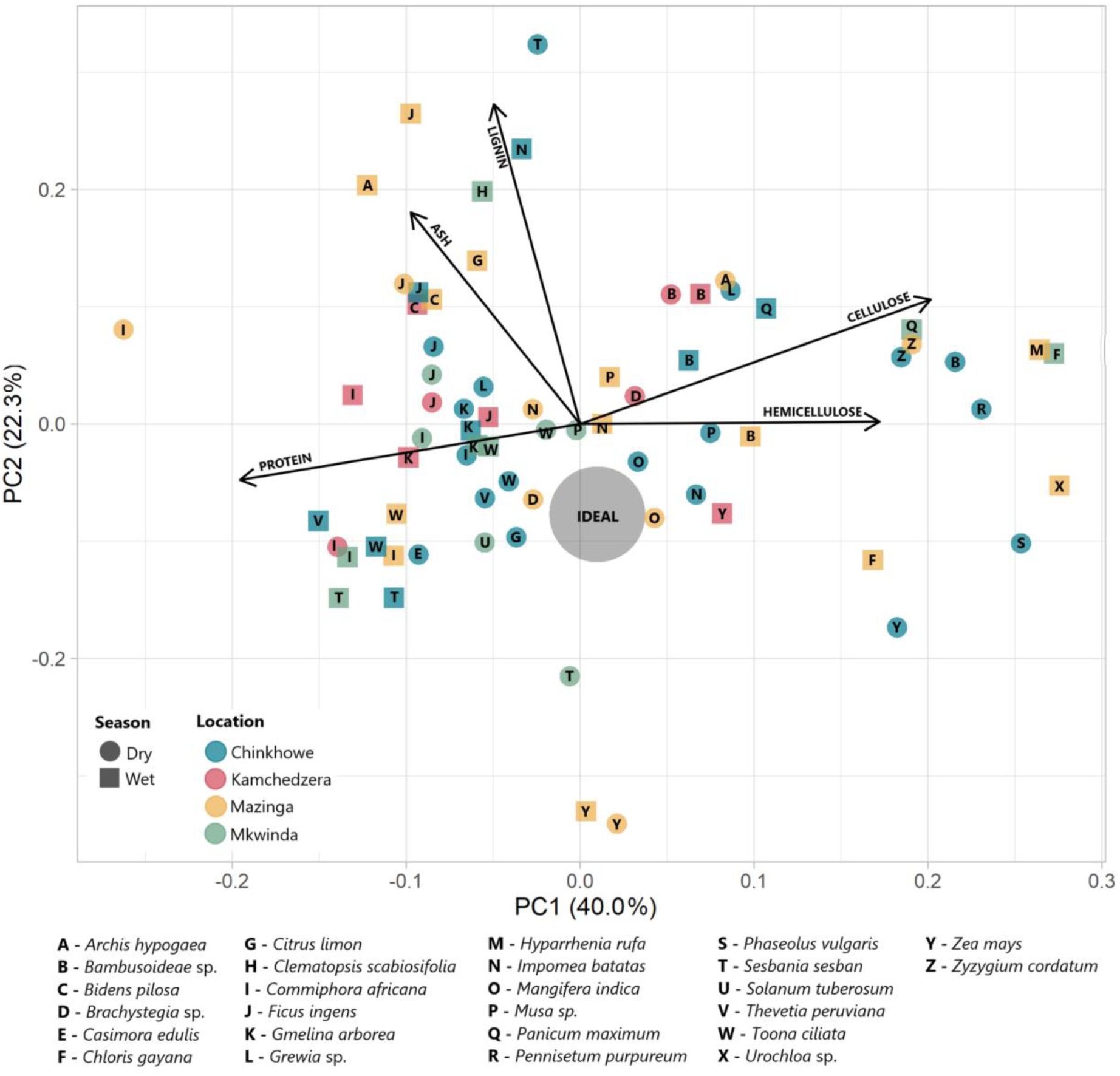
Principal component analysis (PCA) of forages. Each point represents mean nutritional values for one plant species (letter), during one season (shape), at one location (colour). The large grey circle represents an ‘ideal’ nutritional profile based on approximately: ash (9%), protein (15%), lignin (8%), hemicellulose [NDF-ADF] (18%), and cellulose [ADF-ADL] (18%).

Across all sites healthy vegetation cover, as measured by NDVI, was far lower in the dry season than in the wet season: Chinkhowe -53.8%, Kamchedzera -50.6%, Mazinga -59.1%, Mkwinda -54.9% (Figure 4). NDVI peaks around February before steadily declining through to October after which the recovery occurs as new vegetation grows (Figure 4).

**Figure 4.**
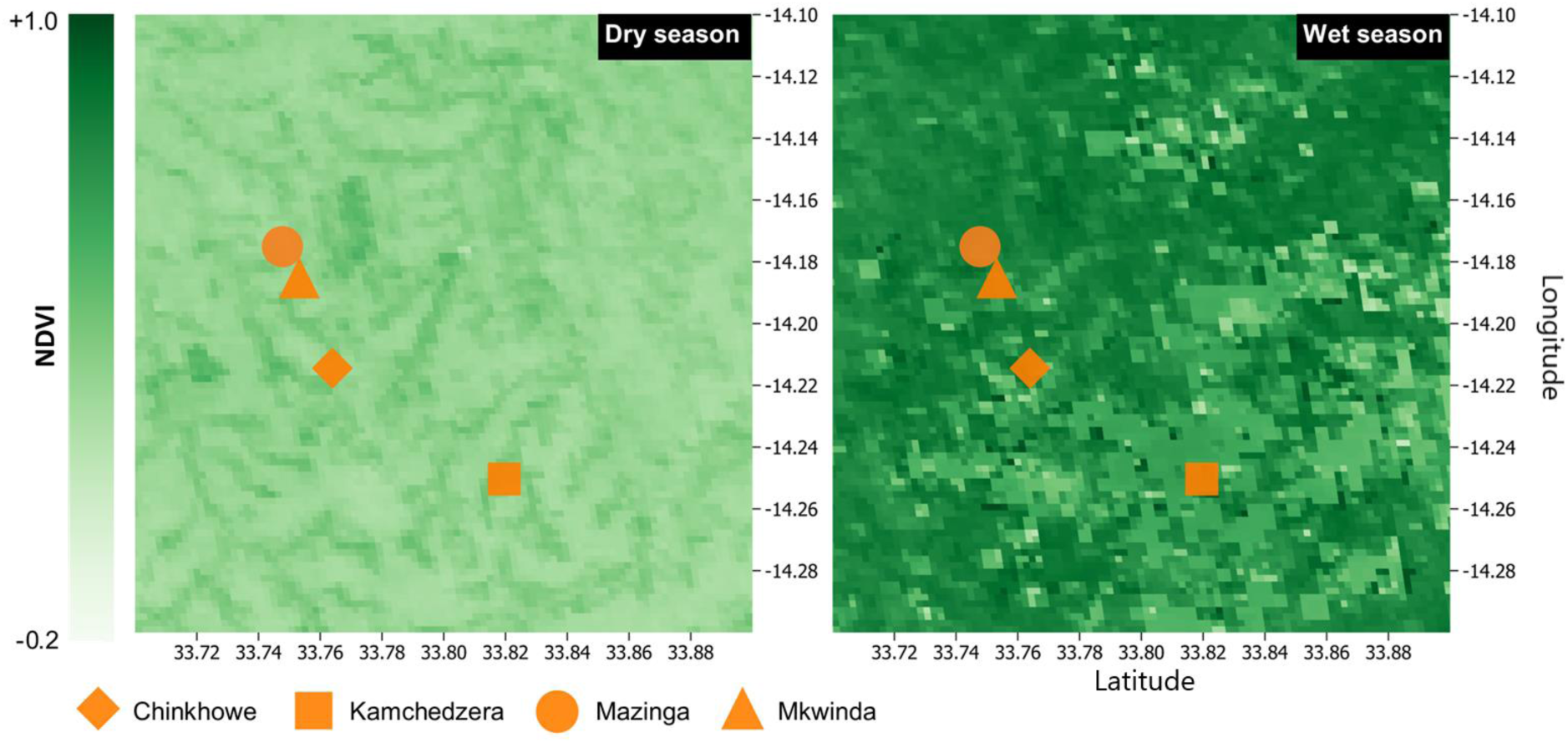
NDVI for the study area during the dry and wet seasons. Data is a 16-day mean NDVI on a 250m grid resolution taken from Moderate Resolution Imaging Spectroradiometer (MODIS) Terra (NASA, 2022b). Dry season data is for the period 12/08/2020 to 27/08/2020 and wet season data for the period 02/02/2021 to 17/02/2021.

**Figure 5.**
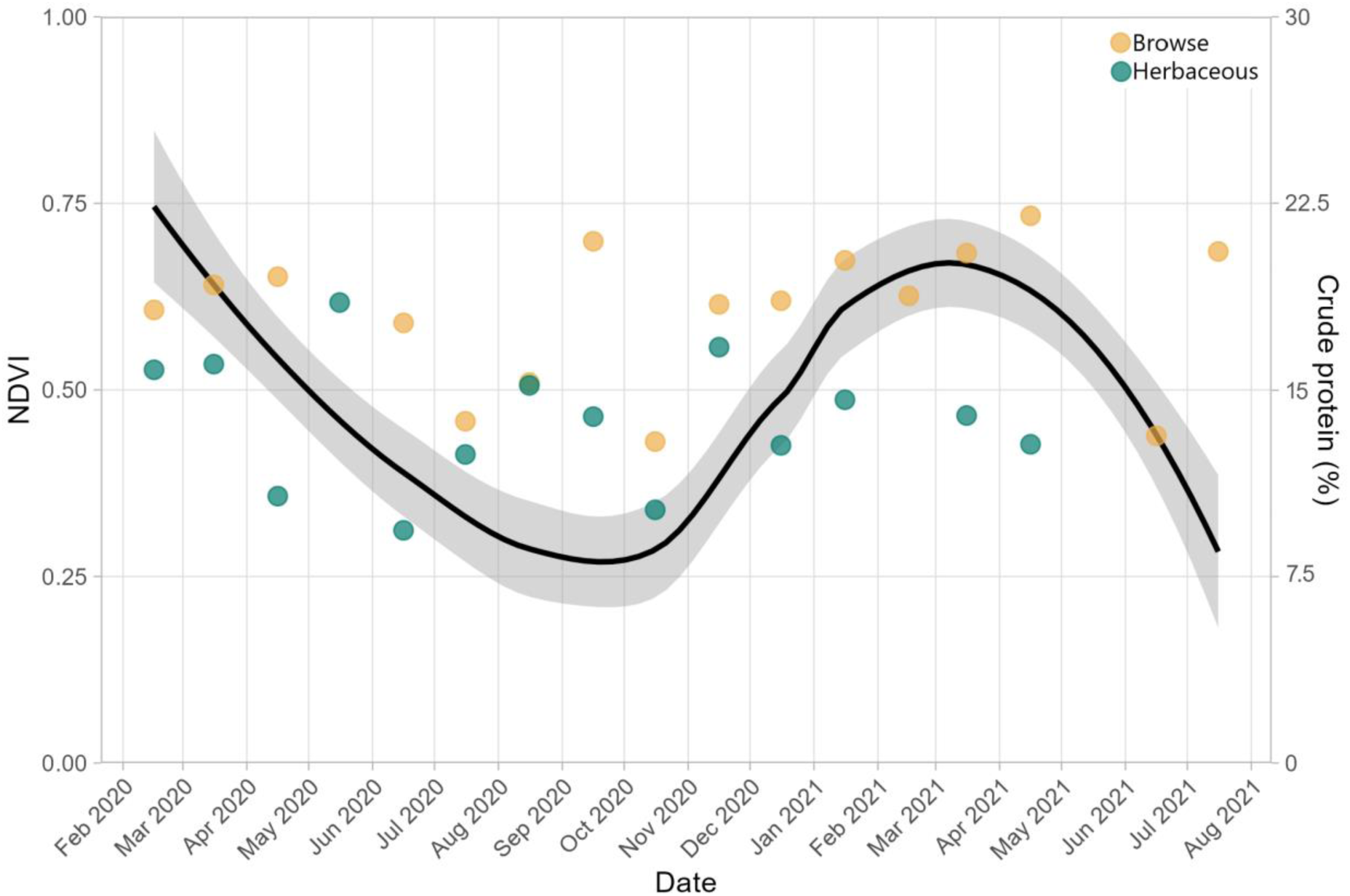
Seasonal fluctuations in NDVI (curve) calculated as the mean NDVI of a 1 km^2^ area at each site. Shaded area represents 95% confidence interval. Overlayed with monthly mean crude protein concentrations (%) of herbaceous and browse species across all sites. NDVI data taken from Moderate Resolution Imaging Spectroradiometer (MODIS) Terra monthly 1km^2^ grid (NASA, 2022b)

## 4 Discussion

Results highlight the extent of the nutritional feed gap that occurs between the wet and the dry season in Malawi and concur with results reported elsewhere in the region (Cooke et al., 2023a; Mphinyane et al., 2015; Naumann et al., 2017). Large reductions in forage availability are exacerbated by reductions in the nutritional value of those forages. The combination of these effects generates a significant reduction in nutritional availability, greater than either measure individually explains. Differences in forage nutrition across sites mean that the exact nature of, and solutions to, the nutritional gap vary location to location.

There was significant variation in the nutritional profiles between plant species, as one may expect. However, whilst browse samples had a higher mean crude protein content (17.4%) compared to herbaceous species (13.8%) (comparable to those seen elsewhere, e.g. Mphinyane et al. [2015]). The browse plant *Commiphora africana* (“African myrrh”) had an especially favourable nutritional profile, being high in CP, but relatively low in ADL. Notably, *C. africana* was present in all four villages across both seasons, with a relatively small reduction in nutritional quality between seasons. At the beginning of the dry season the leaves are desirable to goats, however the plant does defoliate over the course of the season (Le Houerou, 1980; Stuth and Kamau, 1990). Stuth and Kamau (1990) reported that, in Kenya, *C. africana* became an important feed resource for goats during wet season, corresponding to when leaf growth predominantly occurs. Whilst yield was not quantified, its presence at all sites suggests it is well adapted to the region and thus holds potential as a browse for goats. Another browse plant, *Toona ciliata,* also yielded favourable nutritional profiles of high CP and low ADF and ADL. However, was only found at three of the four sites. *Ficus ingens* and *Gmelina arborea* showed notably little variation between sites and seasons, suggesting it may be a reliable contributor to nutrition. Lu et al. (2005) suggest forage ADF concentrations of 23% to be sufficient for growing goats, with anything greater than that potentially inhibiting digestibility. Of the 71 species*seasons collected in this study, 60 yielded ADF concentrations > 23%. Similarly, ADL concentrations were high with 65 species*samples having concentrations greater than the recommended 5%. This suggests the possibility of limited digestibility across available forages, possibly due to the nature plants and their requirements for hardiness to survive in what can be a relatively hostile environment. Goats are however considered “intermediate eaters” and can efficiently consume and digest a wide variety of forages compared to other ruminants (Hofmann, 1989). Whilst PCA coordinates cannot fully quantify the nutritional profiles of plants, they do give a reasonable indication and can be used to provide insight into each location:

**Chinkhowe** - The variation in nutrition, as represented by PCA, was large at this location, though that is partly a consequence of it having the greatest sample size. The browse samples were particularly high in CP, typically exceeding the 15% threshold during both the dry and wet seasons. This is promising in mitigating the seasonal nutritional gap, however that is dependent on sufficient biomass to meet DMI requirements. The wet season nutritional composition of herbaceous species was moderate, though with some examples particularly low in CP (*Bambusoideae* sp. and *Phaseolus vulgaris*). *Mangifera indica* has a nutritional profile especially close to the ‘ideal’.

**Kamchedzera** – Samples from this location had the least variation, though the sample size and species variation were also the smallest of the four locations. No species showed extreme excesses or lack of the major nutrients assessed, something that was observed elsewhere, especially for grasses. Notably, *Zea mays* appeared to be the plant most closely aligned to the ‘ideal’ profile.

**Mazinga –** The largest variation in nutrition was observed at this site which yielded the highest and lowest values on PC1 and the second highest and lowest on PC2. Particularly notable was the low lignin and ash content of *Z. mays* and the high CP content of *C. africana.* Conversely, several plants were particularly low in CP, including *Hyparrhenia rufa* and *Urochloa* sp. though as grasses mainly abundant in the wet season, they could contribute well to DMI so long as CP consumption is met elsewhere in the diet through browse. A number of species, particularly *Brachystegia sp.* and *Grewia* sp. were especially high in ADL. Similar to in Chinkhowe, *Mangifera indica* appeared to have an especially favourable nutritional profile.

**Mkwinda –** Samples from Mkwinda were mostly above CP requirements, as represented by the low PC1 scores. Samples of *Panicum maximum* and *Chloris gayana* were particularly fibrous with low CP contents. Similar to the grasses seen at Mazinga however, they may contribute well to goats DMI, contingent on sufficient CP consumption elsewhere. The herbaceous flowering plant *Musa* sp. had the most favourable nutritional profile, though this was closely followed by *Toona ciliata* and *Solanum tuberosum*.

Significant variations of ADF, and ADL were observed between villages. Differences in ash content were not spatially significantly different, though at p < 0.10 it may be worth exploring in further detail by quantifying the specific minerals in the ash component, especially if that were possible over a greater spatial scale, inclusive of a wider variety of soils. Elsewhere, plant mineral concentrations have been observed to vary across both time and space (Aganga and Mesho, 2008; Mtimuni et al., 1983). Far greater spatial variation may exist at a national level, especially if samples were taken from other soil types (much of Malawi is on lixisol and some of it cambisol soils) or in the north (high altitude, cooler) or south (low altitude, hotter). This highlights the importance of tailoring nutritional interventions to specific circumstances.

The observed seasonal differences in forage CP, with greater concentrations in the wet season compared to dry, has been observed across Sub-Saharan Africa (Naumann et al., 2017; Omphile et al., 2005; Setshogo et al., 2011). Chiphwanya et al. (2017) similarly found reductions in the CP and increased fibre content of grasses in Malawi from the wet to the dry season. This represents a nutritional gap which risks goat health and performance. Whilst CP concentrations appeared to be sufficient for maintenance, with only three examples of plants falling below the 7% maintenance requirement recommended by Pugh (2014), growth may be somewhat limited as they were typically far below optimal levels of 15-17% (NRC, 2006; Salah et al., 2014). This would be especially if DMI was limited through decreased forages availability. Despite nutritional availability appearing preferable in the wet season, there is little evidence to support that goat mortality is reduced at this time. Conversely, elsewhere in Sub-Saharan Africa, goat kid mortality has been found to be greatest during the wet season, possibly due to the higher parasite burden which may be exacerbated by a lower weaning weight caused by inferior nutrition in the dry season (Ameh et al., 2000; Kusina et al., 2000; Zvinorova et al., 2016).

In addition to a decrease in forage quality in the dry season, there is also a reduction in forage quantity. This is evidenced through NDVI which is significantly lower in the dry season compared to the wet. During the dry season the NDVI across most of Malawi, typically < 0.3, representing sparse and barren land. In contrast, during the dry season NDVI is typically > 0.5, a score typically associated with healthy and dense vegetation and its peak productivity. Dry season NDVI scores were 50.6% - 59.1% lower in the dry season than the wet season, which is broadly similar to results observed on the ground elsewhere in the region, for example, Mphinyane et al. (2015) observed a 53.3% reduction in biomass from Autumn to Spring in Botswana. Though forage abundance was not directly measured in this study, the lower proportion of herbaceous samples in dry season is reflective of their reduced abundance as they die back. Meanwhile, browse species, which can tolerate drier conditions, are proportionally more available. If overall forage availability decreases to the point where the animals’ DMI requirements are not met, this could adversely affect animal health and productivity, which would compound the problem of reduced nutritional quality. This may become more extreme as climate change progresses. Across Malawi, remote sensing has provided evidence of decreasing NDVI, decreasing rainfall, and increasing temperatures (Mungai et al., 2020). Future work should integrate nutritional quality measurements with biomass availability, to provide a truer pitcher of total available nutrition, similar to that by Mphinyane et al. (2015) in Botswana.

Seasonal differences in both forage nutrition and availability could be mitigated through the preservation of forages, collected in the wet season for consumption in the dry season. However, the dry season is when goats graze freely, making such interventions less feasible. Additionally, there are significant challenges to forage preservation such as investment, climate, education, and social hurdles (Balehegn et al., 2022). Whilst goat selling/trading does occur year round, there is a peak through July and August (Kaumbata et al., 2020). As this is just after the wet season ends, when nutritional composition and availability is optimal, this provides an opportunity to maximise growth and finishing before sale. However, the typical sale models in Malawi (per head opposed to by liveweight) does not favour this. The use of alternative sale models, such as auctions (ICRISAT, 2022), may be beneficial in addressing this.

### 4.1 Conclusion

The temporal and spatial variations in forage nutrition highlight the dynamic nature of goat husbandry and health in Malawi. The preferential characteristics of the wet season need to be taken advantage of to mitigate for the nutritional-gap that occurs between the dry season and wet season. However, this is easier said than done due to resource limitations and management practices in the region. On the whole, there has been limited forage nutritional work in Malawi and consequently the expansion of the field could help to fill knowledge gaps that can benefit ruminant production and food security in the long-term.

## Acknowledgements

We, the authors, are grateful for the participation of farmers, with whom the study would not have been possible without.

## Author contributions

Cooke, A. S. – Writing, data analysis, laboratory analysis, project management

Mvula, W. – Writing, sample collection, project management

Nalivata, P. - Writing, sample collection, project management

Ventura-Corddero, J. – Writing, data analysis

Gwiriri, L. C. – Writing

Takahashi, T. - Writing, project management, funding acquisition

Morgan, E. R. - Writing, project management, funding acquisition

Lee, M. R. F. – Writing, project management, funding acquisition

Safalaoh, A. - Writing, sample collection, project management

